# ListPred: A predictive ML tool for virulence potential and disinfectant tolerance in *Listeria monocytogenes*

**DOI:** 10.1101/2024.01.29.577690

**Authors:** Alexander Gmeiner, Mirena Ivanova, Rolf Sommer Kaas, Yinghua Xiao, Saria Otani, Pimlapas Leekitcharoenphon

## Abstract

Despite current surveillance and sanitation strategies, foodborne pathogens continue to threaten the food industry and public health. Whole genome sequencing (WGS) has reached an unprecedented resolution to analyse and compare pathogenic bacterial isolates. The increased resolution significantly enhances the possibility of tracing transmission routes and contamination sources of foodborne pathogens. In addition, machine learning (ML) on WGS data has shown promising applications for predicting important microbial traits such as virulence, growth potential, and resistance to antimicrobials. Many regulatory agencies have already adapted WGS and ML methods. However, the food industry hasn’t followed a similarly enthusiastic implementation. Some possible reasons for this might be the lack of computational resources and limited expertise to analyse WGS and ML data and interpret the results. Here, we present ListPred, a ML tool to analyse WGS data of *Listeria monocytogenes*, a very concerning foodborne pathogen. ListPred is able to predict two important bacterial traits, namely virulence potential and disinfectant tolerance, and only requires limited computational resources and practically no bioinformatic expertise, which is essential for a broad application in the food industry.

**AUTHOR SUMMARY:** The contamination of food products with microbes such as pathogenic bacteria is a big concern for the food industry. The rapid detection, characterisation and eradication of microbial contaminants are of utmost importance to ensure safe food products. Fortunately, strict food safety regulations and stringent cleaning protocols prevent the transmission of harmful bacteria to humans. Individual bacteria of the same species can have varying abilities to resist cleaning agents or infect a host, meaning that some pathogen isolates might be more dangerous than others. Novel techniques such as genome sequencing and machine learning can help to determine such differences in individual bacteria. Unfortunately, these techniques require a lot of computational power and expertise that is limited in the food industry. This is why we developed an easy-to-use software tool called ListPred that can be used with few computational requirements and little expertise. ListPred helps food companies to answer two essential questions: how dangerous are *Listeria monocytogenes* pathogens, and how to get rid of them most efficiently?

## INTRODUCTION

*Listeria monocytogenes* is a foodborne pathogen that can cause severe health implications for immunocompromised individuals, the elderly, unborn babies and developing infants [1]. Mortality rates of infected people can even reach between 20-30% [2,3]. However, not all *L. monocytogenes* isolates are equally concerning. Recent research has found differences in niche association and virulence potential on sub-species and even isolate levels [4–7]. As *L. monocytogenes* is mainly transmitted through contaminated food, it is of great concern for the food industry [6,8]. *L. monocytogenes* is ubiquitously distributed in the environment and can enter food processing environments through different channels such as raw produce, equipment, and staff. Once in the production environment, it can manifest itself in different niches and persist over long timespans [9]. Hence, effective sanitation procedures are key in preventing *L. monocytogenes* infections.

Whole genome sequencing (WGS) has found increased popularity in recent years. Many governmental and regulatory agencies, e.g., Denmark, France, and the United States, have already adopted WGS as a standard tool for the surveillance of foodborne pathogens due to its increased genomic resolution. Additionally, the use of bacterial WGS data in combination with ML has shown excellent results for the prediction of important phenotypes such as antibiotic resistance, virulence, and growth potential [10–14]. This highlights the possible benefit of WGS and ML for a more detailed analysis of foodborne pathogens in the food industry. However, the implementation of WGS methods in the food industry has been dragging despite the increasing interest and benefit of possible applications. One of the reasons for this might be limited capacities in a food industry setting. Many food producers do not have the IT infrastructure, computational resources, or bioinformatic expertise required for analysing and interpreting WGS data [15–17]. Consequently, the development of easy-to-use analysis pipelines could facilitate the implementation of WGS methods in the food industry, especially in the early stages.

Here, we introduce ListPred, a quick and easy-to-use ML prediction tool for virulence potential and disinfectant resistance in *L. monocytogenes* at the isolate level. ListPred is accessible through different channels and has three usage variations to accommodate varying user expertise and computational resources. We conducted a small-scale benchmarking study assessing the prediction consistency across different sequencing platforms, i.e., short- and long-read sequencing, and different sequencing data input types, i.e., raw reads or assemblies. In the future, ListPred might be able to aid microbial risk management and sanitation design in the food industry, thereby improving food safety.

## RESULTS

### Comparison of prediction matches across raw and assembled input

For benchmark data set 1, we found a discordance of 8% - 10% when comparing raw read and assembled inputs of both sequencing methods (i.e., short and long reads) and prediction tasks (i.e., virulence potential and disinfectant tolerance). The comparison of predictions for virulence potential from raw and assembled long reads yielded the highest concordance of 92.8%. Predictions of virulence potential and disinfectant resistance comparing raw and assembled short reads showed the lowest concordance of 89.7%. Generally, we observed that using long reads resulted in higher prediction overlaps than short reads when comparing raw reads versus assemblies as input (Table 1, Table S2).

**Table 1.** Comparison of predictions from raw read and assembled sequence data input.

|  | Raw reads vs assembly |  |  |  |  |
| --- | --- | --- | --- | --- | --- |
|  | Benchmarking set 1 |  |  |  | Benchmarking set 2 |
|  | Virulence potential |  | Disinfectant tolerance |  | Virulence potential |
|  | short reads | long reads | short reads | long reads | short reads |
| Match | 87 | 90 | 87 | 88 | 168 |
| Mis-match | 10 | 7 | 10 | 9 | 1 |
| %-match | 89.7 | 92.8 | 89.7 | 90.7 | 99.4 |

For benchmark study 2, we could observe a great overlap between predictions from short-read raw reads and assemblies of 99.4%. This corresponds to only one mismatch where the predicted class was ‘low’ for raw reads and ‘medium’ for assemblies as input (Table 1, Table S2).

### Comparison of prediction matches between short- and long-read sequencing

The comparison of predictions between short- and long-read sequencing for benchmarking data set 1 showed a concordance of 88%-98% for both input types (i.e., raw reads and assemblies) and prediction tasks (Table 2).

**Table 2.** Comparison of predictions from short and long read sequencing platforms.

|  | short vs. long read sequencing |  |  |  |
| --- | --- | --- | --- | --- |
|  | benchmarking set 1 |  |  |  |
|  | virulence potential |  | disinfectant tolerance |  |
|  | raw reads | assembly | raw reads | assembly |
| **Match** | 86 | 92 | 95 | 94 |
| **Mis-match** | 11 | 5 | 2 | 3 |
| **%-match** | 88.7 | 94.9 | 97.9 | 96.9 |

For the prediction of virulence potential, we observed a quite noticeable number of mismatches (12%) between short and long raw reads. On the other hand, comparing short- and long-read assemblies shows a great overlap of predictions of 95% (Table 2, Table S3).

Looking at the prediction of disinfectant tolerance, we observed great concordance of predictions for both raw reads and assemblies. In particular, we found a 98% overlap of predictions for raw reads from short- and long-read platforms and a 97% overlap for assemblies from short- and long-read platforms as input (Table 2, Table S3).

### Comparison of actual and predicted disinfectant tolerance classes

Assessing the concordance of actual versus predicted disinfectant tolerance for benchmark data set 1, the analysis showed marked differences between predictions of different input types. We could see a clear advantage of using assemblies over raw reads with differences in balanced accuracy (BA) scores of up to 14%. More precisely, using short-read assemblies resulted in the highest accuracy with a BA of 0.96, followed by long-read assemblies (BA 0.93), long-read raw reads (BA 0.83), and short-read raw reads (BA 0.82) (Table S4). Comparing the prediction performance across sequencing platforms (i.e., short and long reads), we did not observe as great of a difference as when comparing across sequence input types (i.e., raw reads versus assemblies). In particular, the BA scores for the predictions only differed by 0.01 and 0.03 for raw reads and assemblies, respectively (Table S4).

### Comparison of actual and predicted virulence potential classes

Looking at the concordance of actual and predicted virulence potential for benchmark data set 2, we only saw minor differences comparing the different input types. Using short-read raw reads as inputs resulted in a balanced accuracy (BA) score of 1.0, which indicates perfect agreement. For the assembled input, the comparison resulted in a BA score of 0.99. This BA score corresponds to one misclassification where the isolate is actually in the ‘low’ risk class but predicted by the model as ‘medium’ risk (Table S5).

## DISCUSSION

Short-read sequencing is still the most widely used sequencing method, probably due to the high accuracy of the reads produced, the reasonably low cost, and the availability of third-party provided services. Nevertheless, with advances in long-read sequencing, the accuracy of long-read sequencers continues to improve, and the costs of sequencing in general has been continuously dropping [18,19]. As WGS is slowly gaining more traction in the food industry, it will be necessary for individual producers to assess the best fit for them [20]. Consequently, food producers will most likely use both methods, especially in the current transitional phase to a broader implementation of WGS in the food industry. We conducted a small benchmarking study to assess the effect of short- and long-read sequencing inputs on ListPred predictions. Overall, we observed relatively small differences of maximum 11.3% for predicting virulence potential and 3.1% for disinfectant tolerance. This indicates that ListPred predictions are pretty stable across input from different sequencing platforms.

Genome assembly is a very common step in many sequence analysis workflows. However, assembling genomes requires time, computational resources, and bioinformatics expertise, which might be of limited access in the food industry [17,20]. Hence, we compared the effect of raw reads and assemblies on ListPred predictions. The results of the benchmarking study showed a general difference of about 10%. This was relatively constant for raw read versus assembly comparisons of both short- and long-read sequencing, and for the prediction of virulence potential and disinfectant tolerance. Comparing the actual values with the ListPred predicted values, we could see a clear separation between raw reads and assemblies. The balanced accuracy scores of short and long raw reads were very similar (0.82 and 0.83), as were the short- and long-read assemblies (0.96 and 0.93). This leads us to believe that assemblies as input might lead to better predictions than raw reads. However, we want to highlight that there is a bias in the BA scores. As mentioned in materials and methods, there is a lack of datasets with known virulence potential and disinfectant tolerance phenotypes. This forced us to use isolates for the benchmarking study that were used during model training. Nevertheless, we believe that the bias introduced by this affects the results of different input types equally, making the comparisons still possible. To better understand the prediction ability of ListPred when using different input types, it will be crucial to benchmark the tool with fully independent data sets in the future.

The need for quick and user-friendly analysis tools will increase with the rise of sequencing methods, such as WGS, in the food industry. Initially, there will be a big difference in computational and bioinformatic expertise across food production companies. Bigger producers might have the economic resources for sequencing and computing facilities, whereas medium and smaller-sized producers might struggle to allocate the funding for the implementation of these methods. Hence, we developed ListPred to have a point-and-click web application that alleviates computational resources and can be used with basic computational skills. The output of ListPred is clear and descriptive to enhance interpretability. Microbial data collected by food-producing companies, especially from pathogenic bacteria, is generally confidential, which might limit the willingness to upload data to a web application, even if data is not stored. To ease concerns and facilitate maximum privacy of sensitive data, we additionally distributed ListPred in two stand-alone versions that can be run entirely offline and in-house.

Considering the increasing amount of new data becoming available through a broader implementation of WGS in the food industry, an important consideration might be whether and how the existing models should be updated. In general, there are different approaches to include new data in existing ML models. The most straightforward approach might be to retrain a completely new model of a combined dataset, including the latest data. However, depending on the size of the data set, re-training uses a lot of computational resources and time. There are other methods of including new data in existing models that usually fall into the realm of incremental learning (also called online learning) [21]. The current implementation of ListPred does not accommodate incremental learning approaches. Given that the underlying models of ListPred are trained on relatively small data sets, due to the limited availability of proper data sets, it might be more appropriate to train a new model once new fitting data is available. Nevertheless, to enhance the generalizability of ListPred predictions, the exploration of new data and subsequent updates of the current models should be the topic of future studies.

ListPred is an essential first step towards accessible and quick analysis of WGS data for the food industry. Nevertheless, there are a few yet important limitations that could affect the real-world application of ListPred. Some of these shortcomings, such as the shortage of appropriate data for model training that compromises generalisation ability, the increased disinfectant tolerance through biofilm formation, and the transferability of predictions for pure disinfectant compounds to commercialised products, have been previously discussed in the respective publications of the individual prediction models [22,23].

Other shortcomings are more general to the current barriers of implementing WGS in the food industry. Generally, there is a broad consensus about the possible applications and benefits of WGS methodologies in the food industry. Nevertheless, these methodologies are rather resource intensive and require a lot of expertise, which can become an economic burden. In turn, the benefits of WGS might not necessarily outweigh the additional cost and resources needed, especially for medium and smaller sized food production companies. A possible solution could be to outsource sequencing and analysis to third-party providers. However, this most likely increases the result turnaround times, which limits applicability for time-sensitive situations.

Another important barrier for WGS application in the food industry is the current lack of regulatory frameworks and guidelines regarding these methods [16,24]. Without a clear regulatory system, food industry stakeholders might fear possible repercussions due to improved tracing of contamination and foodborne illness sources. The combined effort of regulatory food safety agencies and food industry representatives to tackle this barrier will be crucial for a broader implementation of WGS methods in the food industry and the creation of novel structures to fight the spread of foodborne diseases.

In conclusion, ListPred aims to enable quick and accessible WGS analysis in the food industry. The increasing implementation of sequencing technologies in the food industry will raise the need for easy-to-use tools that don’t require computational or bioinformatic expertise. The phenotypic traits predicted by ListPred could give risk managers important insight into the potential threat of *L. monocytogenes* isolates found in the production environment and how to eliminate them more effectively. This information might even be considered in the design of risk mitigation strategies and sanitation plans. Apart from this, ListPred is an important example of possible applications and possible benefits of WGS implementation in the food industry. Considering further application showcases, lowering sequencing prices, and the development of novel end-to-end analysis tools might shift current cost-benefit evaluations in favour of a more routine implementation of WGS in the food industry.

## MATERIALS AND METHODS

### Design and Implementation

The predictive bioinformatics pipeline ListPred is realised through the Snakemake workflow tool [25]. The pipeline combines whole genome sequencing data pre-processing, machine learning predictions, and easy interpretable reporting. ListPred can process various inputs, namely raw read or assembled sequencing data from short- or long-read sequencing experiments. The sequencing data is pre-processed according to the input type (Fig. 1). First step in the pipeline is screening the sequence data for the presence of reference gene features used for the ML prediction. This is done for both sets of reference gene features used to predict virulence potential and disinfectant resistance. The screening yields alignment percent identity scores for each reference gene feature and is realised using tblastx [26] for assemblies and KMA [27] for raw reads. All the results from the individual screenings are gathered in two matrices used as input for ML, i.e., one for virulence potential and one for disinfectant tolerance prediction. The ML models used for prediction in this pipeline were developed in previous studies [22,23]. The prediction results are reported in tabular format and are either low, medium, or high for virulence potential and sensitive or tolerant for disinfectant tolerance prediction.

**Fig. 1.**
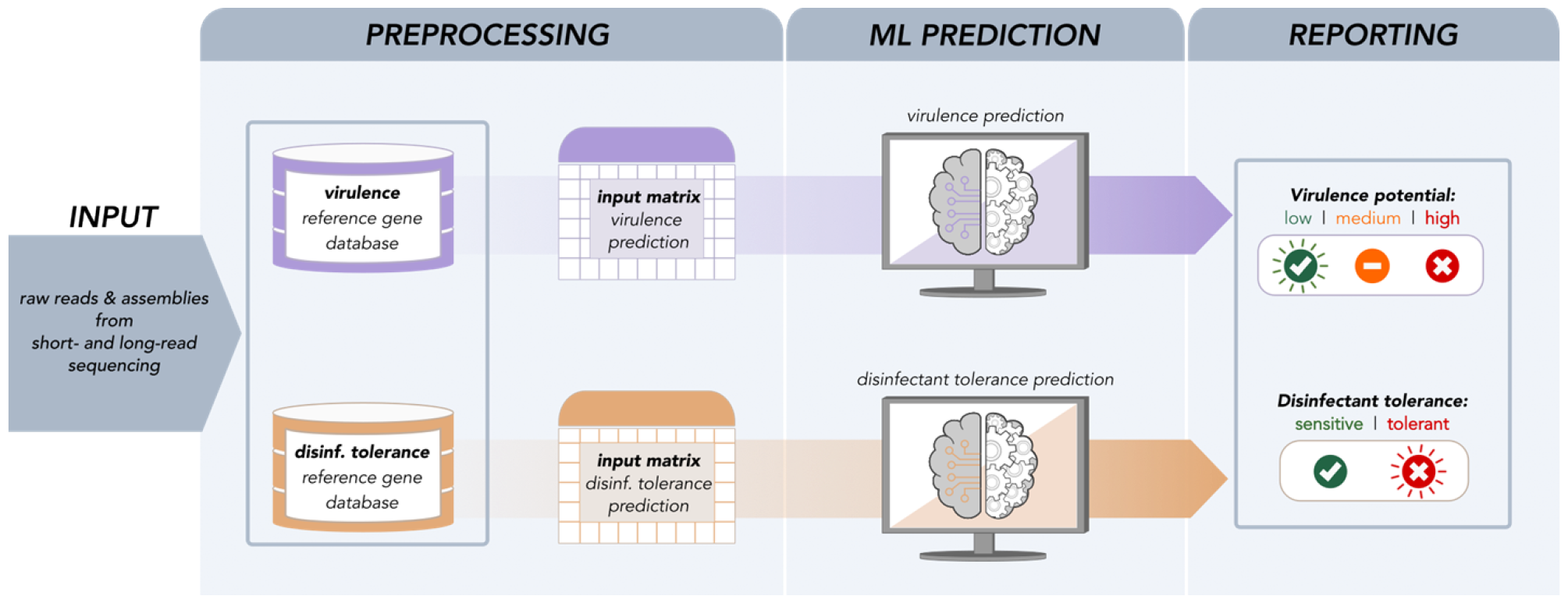
Schematic overview of ListPred workflow. This is a schematic overview of the three parts, i.e., preprocessing, ML prediction, and reporting, that are part of the ListPred workflow.

### Distribution

ListPred is available in three variations depending on user experience and computational prerequisites. For users that are comfortable with the execution of command line programs, look to process multiple isolates at the same time, and run the analysis offline, the source code for the ListPred Snakemake pipeline is available on GitHub (https://github.com/genomicepidemiology/ListPred/tree/master/workflow).

For users with command-line experience who are looking to process multiple isolates and want a low-effort execution, ListPred is available as Docker Image on Docker Hub (https://hub.docker.com/r/genomicepidemiology/listpred). This solution can be run offline by manually downloading the Image to a local server or online (i.e., Docker handles the download from Docker Hub and subsequent execution).

For users with limited computational experience, a point & click solution in the form of a web application is hosted by the Center for Genomic Epidemiology at the Technical University of Denmark (DTU) and can be accessed by the following link (http://genepi.food.dtu.dk/listpred)

### Benchmarking

To assess the concordance of prediction results regarding different sequencing platforms (i.e., short-read and long-read sequencing) and different sequencing input types (i.e., raw reads and assemblies), we performed a small-scale benchmarking case study for ListPred using two data sets. Finding datasets that share WGS data and describe virulence potential or disinfectant phenotypes that are comparable to the prediction output of ListPred proved to be quite difficult. This is why we had to resort to using the same sequencing data that was used to develop the predictive models, being well aware that this introduced some bias in the evaluation. Nevertheless, since the main aim of this benchmarking case study is to assess differences between input types rather than evaluate the performance of ListPred, this is of less concern. The bias introduced by using isolate data that the ML models have previously seen affects all predictions similarly. In theory, this should still enable us to assess some general patterns when comparing the differences in prediction. Due to the difficulty of finding appropriate benchmarking datasets, we do not have the actual phenotypes describing the ground truth for all isolates. However, we do have actual disinfectant tolerance phenotypes for data set 1 and virulence potential phenotypes for the benchmarking data set 2.

### Benchmarking data set 1

The *L. monocytogenes* isolates for the benchmarking data set 1 (n=97) were selected from Ivanova et al. (2023) [28] and cover a broad range of geographical regions, isolation sources and isolate sub-types. The isolates’ short-read WGS data was acquired from the European Nucleotide Archive (ENA). The short-read sequencing data was assembled by trimming the raw reads with bbduk2 from BBTools (v36.49) [30] and performing de-novo assembly using SPAdes (v3.11.0) [31]. We also excluded contigs of 500bp length and smaller. In addition, we performed long-read sequencing for the selected isolates. For this, isolates were grown on trypticase soy agar plates at 37°C overnight. Total DNA was extracted using the Quick-DNA HMW Magbead Kit (Nordic BioSite, Denmark) with minor modifications, i.e., increased starting material (half 10-μL loop) and beads volume (50 μL) and all mixing steps (including those with beads) were done using a vortexer at 1400 rpm instead of a rotator. ONT libraries were prepared using ligation sequencing kit SQK-LSK109 with native barcoding expansion kits EXPNBD104 and EXP-NBD114 (Oxford Nanopore Technologies, Oxford, UK), loaded onto R9.4.1 flow cells (Oxford Nanopore Technologies) and sequenced using GridION (Oxford Nanopore Technologies) for 72 h. The Nanopore reads were assembled using Flye (v2.9.1) [29] with the ‘--nano-hq’ flag. Lastly, we extracted the laboratory-tested disinfectant tolerance phenotypes, i.e., sensitive or tolerant, from the metadata of the study. A detailed description of the isolates, including ENA accession numbers, metadata, and phenotypes, can be found in Table S1.

### Benchmarking data set 2

The information for benchmarking data set 2 was collected from Gmeiner et al. (2024) [23] and consisted of short-read WGS from 169 *L. monocytogenes* isolates. The isolates described in that study were collected through national surveillance programs in France and Denmark and covered the public health sector and the food industry. We further extracted the isolates’ corresponding phenotypes, i.e., low, medium, or high virulence potential described in the study. The corresponding assemblies were constructed using the same workflow as described for benchmarking data set 1. For a detailed description of the isolates’ accession numbers and phenotypes, see Table S1.

## ACKNOWLEDGMENTS

This work was supported by Karl Pedersen og Hustrus Industrifond (DI-2019-07020), the Danish Dairy Research Foundation, and the Milk Levy Fund.

## CONFLICT OF INTEREST

The authors of this publication do not have any conflicts of interest to declare.

## SUPPORTING INFORMATION

See attached file.

## DATA AVAILABILITY

All WGS sequencing data is publicly available on ENA (see accession numbers in Table S1). The source code of the ListPred pipeline can found on GitHub (https://github.com/genomicepidemiology/ListPred/tree/master/workflow).

## Notes

### Competing Interest Statement

The authors have declared no competing interest.

